## Supplementary Data S1 for "Genome of the endangered eastern quoll (*Dasyurus viverrinus*) reveals signatures of historical decline and pelage color evolution"

**Supplementary Data S1: Alignment of dasyuromorph ASIP orthologs**

Mfas – Numbat (*Myrmecobius fasciatus*)

Tcyn – Thylacine (*Thylacinus cynocephalus*)

Astu – Brown antechinus (*Antechinus stuartii*)

Afla – Yellow-footed antechinus (*Antechinus flavipes*)

Dviv – Eastern quoll (*Dasyurus viverrinus*)

Shar – Tasmanian devil (*Sarcophilus harrisii*)

Missing sequence in the Tasmanian devil

```
Alignment Of ASIP protein - MAFFT (v7.511)

Mfas       MTAKHLFLPFLLACLWFLAAYCHLAEEEKWSKDKGLGRNSMNLTDFPSVSIVALNKKSKK
Tcyn       MTAKHLLLPFLLVCLWFLAAYCHLAEEEKWSKDRSLGRNSMNLPDFPSVSIVALNKKSKK
Astu       MTAKHLFLPFLLACLWFLAAYCHLAEEEKWSKDRGLGRSSMNLPDFPSVSIVALNKKSKK
Afla       MTAKHLFLPFLLACLWFLAAYCHLAEEEKWSKDRSLGRSSMNLPDFPSVSIVALNKKSKK
Dviv       MTAKHLFLPFLLACLWFLAAYCHLAEEEKWSKDRGLGRSSMNLPDFPSVSIVALNKKSKK
Shar       -----------------------------------------------------LNKKSKK
                                                                *******

Mfas       SIRKEIETKKSSEKKAVVKKSPSRSNCAATGAFCQPHTLSCCEPCASCYCRFFGRVCSCR
Tcyn       SIRKEIETKKSSEKKAVVKKSSSGSNCAATGAFCQPQTLSCCNRCATCHCRFFRSSCSCR
Astu       SIRKEIETKKSSEKKAVVKKSSSASTCAATGAFCQPQTISCCDKCDTCHCRFFGSVCFCR
Afla       SIRKEIETKKSSEKKAVVKKSSSASTCAATGAFCQPQTISCCDKCDTCHCRFFGSVCFCR
Dviv       NIRKEIETKKSSEKKAVVKKSSSASTCAATGAFCQPQTISCCNKCDTCHCRFFGSVCFCR
Shar       NIRKEIETKKSSEKKPVVK--LSASTCAATGAFCQPQTISCCNKCDTCHCRFFGSVCFCR
           .**************.***   * *.**********:*:***: * :*:****   * **

Mfas       LFQQSC
Tcyn       LFQPGC
Astu       PFMRKC
Afla       PFMRKC
Dviv       QFLRKC
Shar       PFLRKC
            *   *
```
