## Supplementary Data S2 for "Genome of the endangered eastern quoll (*Dasyurus viverrinus*) reveals signatures of historical decline and pelage color evolution"

**Supplementary Data S2: Alignment of *ASIP* exon 1 region**

Shar 1 - mSarHar1.11 (Region = NC_045427.1:421297207-421297632)

Shar 2 - SarHar_Dovetail_2.0 (Region = VOSF01009762.1:640079-640504)

Afla - AdamAnt_v2 (Region = NC_067399.1:439018275-439020331)

Dviv - UniMelb_DasViv_v1.0 (Region = CM036998.1:421733752-421736969)

*ASIP* exon 1

Putative eastern quoll insertion boundaries (10bp highlighted)

Putative Tasmanian devil deletion boundaries (10 bp highlighted)

```
Alignment Of ASIP Exon 1 Region - MAFFT (v7.511)

Shar1      tcttacccatcttttaatgggtattaaataccgaattctctag-aagccttctttgaaac
Shar2      tcttacccatcttttaatgggtattaaataccgaattctctag-aagccttctttgaaac
Dviv       tcttacccatcttttaatgggtattaaataccgaattctctagaaagccttctttgaaac
Afla       tcttacccatcttttaatgaatattaaatactgaatcctctagaaagccttctttgaaac
           *******************..**********.****.****** ****************

Shar1      agtctccctttacactgtccatatcttgtattttcataattatctgcaagttgcctcctc
Shar2      agtctccctttacactgtccatatcttgtattttcataattatctgcaagttgcctcctc
Dviv       attctccctttacactgtccctatcttgtattttcataattatctacaagttgcctcctc
Afla       agtctccctttacactgttcatatcttgtattttcataattatctgcaagttgcctcctc
           * ****************.* ************************.**************

Shar1      cattagaaggtgaattccttgagagtaggaaattaggctttttcctttccttaaaacccc
Shar2      cattagaaggtgaattccttgagagtaggaaattaggctttttcctttccttaaaacccc
Dviv       cattagaaggtgaattccttgagagtaggaaa-taggctttttcctttccttaaaacccc
Afla       tattagaacatgaattccttgagagtaggaaattaggctttttcgtttccttaaattccc
           .******* .********************** *********** ********** .***

Shar1      acggttaagggcagtggctg----------------------------------------
Shar2      acggttaagggcagtggctg----------------------------------------
Dviv       agggttaagggcagtgtgtgtgtgtatatatatatatatatatatatatatacatacaca
Afla       agggataagggcagtggctg----------------------------------------
           * ** ***********  **

Shar1      -----gcacatagtatcaatcaatcaat--------------------------------
Shar2      -----gcacatagtatcaatcaatcaat--------------------------------
Dviv       cacacacacatagtatcaatcaatcaataaacatttattgtgtctacacacctgtcagat
Afla       -----gcacatagtatcaatcaatcaat--------------------------------
                .**********************


Shar1      ------------------------------------------------------------
Shar2      ------------------------------------------------------------
Dviv       tggctaagatgacaggaaaaaataataatgattgttggaggggatgtgggaaaactggga
Afla       ------------------------------------------------------------


Shar1      ------------------------------------------------------------
Shar2      ------------------------------------------------------------
Dviv       cattgttgcattgttggtggagttgtgaacgaatccaaccattttggagagttgtttgga
Afla       ------------------------------------------------------------


Shar1      ------------------------------------------------------------
Shar2      ------------------------------------------------------------
Dviv       actatgctcaaaaagttatcaaactgtgcataccctttgatccagcagtgttactactgg
Afla       ------------------------------------------------------------


Shar1      ------------------------------------------------------------
Shar2      ------------------------------------------------------------
Dviv       gcttatatcccaaagagattataaagcagggaaagggacctgtatgtgcacgaatgtttg
Afla       ------------------------------------------------------------


Shar1      ------------------------------------------------------------
Shar2      ------------------------------------------------------------
Dviv       tggcagccctttttgtagtggctagaaactggaagctgaatggatgcccatcagttggag
Afla       ------------------------------------------------------------


Shar1      ------------------------------------------------------------
Shar2      ------------------------------------------------------------
Dviv       aatggctgaataaattgtggtatatgaatactatggaatattactgttctgtaagaaatg
Afla       ------------------------------------------------------------


Shar1      ------------------------------------------------------------
Shar2      ------------------------------------------------------------
Dviv       accaacaggatgatttcagaaaggcctggagagacttacatgaactgatgctgagtgaaa
Afla       ------------------------------------------------------------


Shar1      ------------------------------------------------------------
Shar2      ------------------------------------------------------------
Dviv       tgagcaggaccaggagaacattatatacttcaacaacaatactatatgatgcccagttct
Afla       ------------------------------------------------------------

Shar1      ------------------------------------------------------------
Shar2      ------------------------------------------------------------
Dviv       gatggacctggccatcctcagcaacgagatcaaccaaatcatttccaatggagcagtaat
Afla       ------------------------------------------------------------


Shar1      ------------------------------------------------------------
Shar2      ------------------------------------------------------------
Dviv       gaactgaaccagctatgcctagagaaagaactttgggagatgacgaaaaaccaatacatt
Afla       ------------------------------------------------------------


Shar1      ------------------------------------------------------------
Shar2      ------------------------------------------------------------
Dviv       gaattcccaatccctatatttatgcccacctgcatatttgatttcctccacaagctaatt
Afla       ------------------------------------------------------------


Shar1      ------------------------------------------------------------
Shar2      ------------------------------------------------------------
Dviv       gcacaatatttcagaatcagattctttttatacagcaaaatatgttttggtcatgaatac
Afla       ------------------------------------------------------------


Shar1      ------------------------------------------------------------
Shar2      ------------------------------------------------------------
Dviv       ttattgtatatctaatttatattttaatgtatttaacatctactggtcatcctgccatct
Afla       ------------------------------------------------------------


Shar1      ------------------------------------------------------------
Shar2      ------------------------------------------------------------
Dviv       aggggaaggggtggggggtgggaggcgaaaaattggaacaagaaagttggcaattgttaa
Afla       ------------------------------------------------------------


Shar1      ------------------------------------------------------------
Shar2      ------------------------------------------------------------
Dviv       tgctgtaaagttatccatgcatataacctgtaaataaaaggctattattaaaaaaaaatt
Afla       ------------------------------------------------------------


Shar1      -----------------------aaacatttattgtgtctattaaagtaagcaattcatc
Shar2      -----------------------aaacatttattgtgtctattaaagtaagcaattcatc
Dviv       tttttttaaattaaaaaaaaaaaaaacatttattgtgtctattaaagcaagcaattcatc
Afla       -----------------------aaacatttattgtgtctattagagcaagcaattcatc
                                  *********************.**.************

Shar1      aatgcttaatgcccagtgactaactgcccctcctcacaccttcca---------------
Shar2      aatgcttaatgcccagtgactaactgcccctcctcacaccttcca---------------
Dviv       aatgcttaatgccccgtgactaactgcccctcctcacaccttccacagtggtttatttat
Afla       aatgcttaatgccc-atgactgactgcccctcctcacaccttccacaggggtttatttat
           ************** .*****.***********************

Shar1      ------------------------------------------------------------
Shar2      ------------------------------------------------------------
Dviv       tttggttctttttaatgcacatattgtgtgatagttgtgggtaccacatctcccttctag
Afla       tttcgttcttcttaatgctcatattgtgtgatagttgtgggtaccacatctaccttctag


Shar1      ------------------------------------------------------------
Shar2      ------------------------------------------------------------
Dviv       aacatgtgctccctacagatgggattcaggaggtatttaataaatgttcattgtatttgt
Afla       aatatgtgctccctacagatgggattcaggaggtatttaataaatgttcattgtatttgt


Shar1      ------------------------------------------------------------
Shar2      ------------------------------------------------------------
Dviv       tgttctgacatagccagaccgtaacagtaaatgctgctaagcagaggccatccttctgtt
Afla       tgctttgacattgccaggccatagcagtaaatgctgctaagcagaggccatccttctgtt


Shar1      ------------------------------------------------------------
Shar2      ------------------------------------------------------------
Dviv       ttcctccagtcctcccagtgtcttctcatctccccacagtttttgtctttttcctttcct
Afla       ttcctccagtcctcccaacatcttctcttctccccacaatttttgtctttttcctttcct


Shar1      ------------------------------------------------------------
Shar2      ------------------------------------------------------------
Dviv       cccactgcacttgtctcccctcccctgctttagggctagaccactctgtggctcatgggg
Afla       cccactgcacttgtctcccctctcctgctttagggctagaccactctgtggctcatgggg


Shar1      ------------------------------------------------------------
Shar2      ------------------------------------------------------------
Dviv       tcaccatccagggcagatctcccccactcccccttctccccagccttcacacatcccaat
Afla       tcaccatccaaggcagatctcccccacttccccttctccccagccctcacacatcccaat

Shar1      ------------------------------------------------------------
Shar2      ------------------------------------------------------------
Dviv       cccatcttgacttctggttttctgttctgcttagaagttccaggatgacagctaagcatc
Afla       cccatcttgacttctggttttctcttctgcttagaagttccaggatgacagctaagcatc


Shar1      ------------------------------------------------------------
Shar2      ------------------------------------------------------------
Dviv       tgttccttcccttccttctggcctgcctgtggttcctggctgcctactgccacctggctg
Afla       tgttccttcccttccttctggcctgcctgtggttcctggctgcctactgccacctggctg


Shar1      ------------------------------------------------------------
Shar2      ------------------------------------------------------------
Dviv       aggaagagaaatggagtaaggataggggtctgggaagaagctccatgaacctgcctgact
Afla       aggaagagaaatggagtaaggataggagtctgggaagaagctccatgaacctgcctgact


Shar1      ------------------------------------------------------------
Shar2      ------------------------------------------------------------
Dviv       ttccttctgtgtccatcgtgggtgagtagcctggccagccactatctctgacacaggact
Afla       ttccttctgtgtccatcgtgggtgagtagcctgaccagccaccatctctaacacaggact


Shar1      ------------------------------------------------------------
Shar2      ------------------------------------------------------------
Dviv       agtgtgcagggcgggcctatgatctctatcct----------------gggcttgggctt
Afla       agagtgcagggggggcctatgatcttcctcctgctcccttagcctagagggcttgggctt


Shar1      ------------------------------------------------------------
Shar2      ------------------------------------------------------------
Dviv       cgttttccccttttgggagattccgggtggcctccctcaaatgcagctacctctctcaag
Afla       catttcccccttctgggagattctcggtggcctccctcaaatgcagctacctctctcaag


Shar1      ------------------------------------------------------------
Shar2      ------------------------------------------------------------
Dviv       ttctagctattcttgatagtggtgctcactccccatttagtctatagattgggaaccaca
Afla       ttctagctgttcttgatagtggcactcacttcccatttagtctgtagattaggaactaca


Shar1      ------------------------------------------------------------
Shar2      ------------------------------------------------------------
Dviv       agaactgggttccaatacctgagaaaatcacttctgaaacccactctcagcatttcagat
Afla       agaactgggttccaatacctgagaaaatcacttctgaaacccactctcagcatttcaggt


Shar1      ------------------------------------------------------------
Shar2      ------------------------------------------------------------
Dviv       ttcttcttattaaaaatggggaggggggatttaaatgctctctaatgtcctttccagatc
Afla       tccttctttattaaaatggggaggggggatttaaatgctctctaaagtcccttccagatc


Shar1      ------------------------------------------------------------
Shar2      ------------------------------------------------------------
Dviv       taaaaattttatgatcccttctactaaatggctacctcccctgccttttcttcagctgct
Afla       taaaaatgttatggttccttctactaaatggctacctcccctgccttttcttcaaccgct


Shar1      ------------------------------------------------------------
Shar2      ------------------------------------------------------------
Dviv       tgccagcctgaagtccctatggagtcacagagccttattaatcaaatctggagaaaagtc
Afla       tatcagcctgaagtccctatggagtcac--agccttattaatcaaatctggagaaaagtc


Shar1      ------------------------------------------------------------
Shar2      ------------------------------------------------------------
Dviv       agatgaatgtaactcctttctatagctaatgttcagcatatgagaggtgagctttaatga
Afla       agatgaatgtaactcctttcaatagccaatgttcagtatatgagaggtgggctttaatga


Shar1      ------------------------------------------------------------
Shar2      ------------------------------------------------------------
Dviv       gacctttgattctacctgaaaaacagggtctagaccaatgagatttcagtgcatctaata
Afla       gacttttgattctacctgaaaaacagggtctagatcaatgagatttcagtgcatctaata


Shar1      ------------------------------------------------------------
Shar2      ------------------------------------------------------------
Dviv       tagattaaatgtgaaattcttctaaacatatttccacatttatcattctgcataagaaaa
Afla       tagattaaatgtgaaattcttctaaacgtatttccacatttatcattttacacaagaaaa


Shar1      ------------------------------------------------------------
Shar2      ------------------------------------------------------------
Dviv       atcagatcaaaaagggggg--gaaatgagaaagaaaaaagcaagcaagcaacaaaaatga
Afla       atcagatcaaaaagggggggagaaataagaaagaaaaaagcaagcaagcaacaaagatga


Shar1      ------------------------------------------------------------
Shar2      ------------------------------------------------------------
Dviv       gggagaagcgagatggaaagcaattttattttgatacaattgacttcctttgcaaatctt
Afla       gggagaagcaaggtggaaagcaattttatttagatacaattggcttcctttgcaaatc-t


Shar1      ------------------------------------------------------------
Shar2      ------------------------------------------------------------
Dviv       ttttttttcatgagttgaaaaacattgtgctggaaagggtttacaggcttcggcccagac
Afla       ttttttttcatgagttgaaaaacattatgctggaaagggtttataggtttccccccagag


Shar1      ------------------------------------------------------------
Shar2      ------------------------------------------------------------
Dviv       tccaaaagggccccatggcaaaaaaaaagttaagagcctatctagatagtgctggagact
Afla       tcccaaagggccccatggc--aaaaaaagttaagagcctatctagatagtgctggagact


Shar1      ------------------------------------------------------------
Shar2      ------------------------------------------------------------
Dviv       gagggattaggaaaggtagtctctaagactagtttaaaggtttccacaaacctcttggtc
Afla       aagggattaggaaaggtggtctccaagactagttttgaggtttccacaaacctcttggtc


Shar1      ------------------------------------------------------------
Shar2      ------------------------------------------------------------
Dviv       tcaactgttcccagaacccaaggaaaaaaagggggcaatt---gaaactccttttgaccc
Afla       tcaactgttccaagaacccaaggaaaaaaagggggcaattaaagaaactccctttg----


Shar1      ------------------------------------------------------------
Shar2      ------------------------------------------------------------
Dviv       tgagatactaggaggggaatgccaaaagtggacaagaatcaaatttcttttctgtctagc
Afla       ------------------------------------------------------------


Shar1      ------------------------------------------------------------
Shar2      ------------------------------------------------------------
Dviv       ctgtaagacttcctctattctcaggctttctcctctttgtttgtgaggcacttacctcaa
Afla       ------------------------------------------------------------


Shar1      ------------------------------------------------------------
Shar2      ------------------------------------------------------------
Dviv       tccctggcctttgctcttttggctcctctagcattgaaatggtcaggcaaaaatggtcag
Afla       ------------------------------------------------aaaaaaagtc--


Shar1      ----------------------------------------------------------ca
Shar2      ----------------------------------------------------------ca
Dviv       gggtctccactcctgcctccgggagacccccttcccagaggtctccctcttcccaattca
Afla       ---tctccacccctccctccgggagactcccttcccagaggcctccctcttcccaattca
                                                                     **

Shar1      gtctatccctggccctgccagtcagacatatcttctcttcattttgttttctttaaagca
Shar2      gtctatccctggccctgccagtcagacatatcttctcttcattttgttttctttaaagca
Dviv       gtctatccctggccctgccagtcagacatatcttcttttcattttgttttctttgaagca
Afla       gcctatccctggccctgccagtcagacatatcttctcttcattttgttttctttgaagca
           *.**********************************.*****************.*****

Shar1      ctgaacaaaaaatccaagaagaacatcaggaaagagatagaaaccaagaaatcttccgag
Shar2      ctgaacaaaaaatccaagaagaacatcaggaaagagatagaaaccaagaaatcttccgag
Dviv       ctgaacaaaaaatccaagaagaacatcaggaaagagatagaaaccaagaaatcctccgag
Afla       ctgaacaaaaaatccaagaagagcatcaggaaagagatagaaaccaagaaatcctccgag
           **********************.******************************.******
```
